# Predicting Cellular Drug Sensitivity using Conditional Modulation of Gene Expression

**DOI:** 10.1101/2021.03.15.435529

**Authors:** Will Connell, Michael Keiser

## Abstract

Selecting drugs most effective against a tumor’s specific transcriptional signature is an important challenge in precision medicine. To assess oncogenic therapy options, cancer cell lines are dosed with drugs that can differentially impact cellular viability. Here we show that basal gene expression patterns can be conditioned by learned small molecule structure to better predict cellular drug sensitivity, achieving an *R^2^* of 0.7190±0.0098 (a 5.61% gain). We find that 1) transforming gene expression values by learned small molecule representations outperforms raw feature concatenation, 2) small molecule structural features meaningfully contribute to learned representations, and 3) an affine transformation best integrates these representations. We analyze conditioning parameters to determine how small molecule representations modulate gene expression embeddings. This ongoing work formalizes *in silico* cellular screening as a conditional task in precision oncology applications that can improve drug selection for cancer treatment.

## 1 Introduction

Ideally, personalized tumor profiling could be used to predict an individual patient’s clinical response to a chemotherapeutic agent. The abundance of omics information fuels the promise of such data-driven personalized medicine [14]. Although many approaches attempt to model tumor drug response as a function of cellular features, this remains an open challenge. Previous work focuses on predicting summary metrics of drug sensitivity, such as inhibitory concentration of a drug at 50% cellular viability (IC50) or area under the dose-response curve (AUC). To maximize the therapeutic window of cancer chemotherapeutics, which can have severe side effects, dosage choice is also a key component of precision oncology efforts.

Publicly-available characterizations of cancer model systems such as immortalized cell lines facilitate development of pharmacogenomic models to guide personalized treatment. Many machine learning methods operate under the hypothesis that tumor molecular state determines drug sensitivity. We focus on modeling cellular drug sensitivity as a function of gene expression patterns that depends on small molecule structure and dosage. We hypothesize that models with stronger relational inductive biases, defined by a conditional formulation and expressed by architectural assumptions, will outperform a naive modeling approach. Neural networks are well suited for conditional model formulation due to their architectural flexibility, proven success integrating diverse data types, and most importantly, ability to learn hierarchical feature representations [4, 7].

To our knowledge, previous work modeling cellular sensitivity has only explored integration of different data types as features via concatenation [4, 2]. Given the broad success of conditioning inputs by learned feature representations in visual question-answering, style transfer, generative Learning Meaningful Representations of Life Workshop at NeurIPS 2020. https://www.lmrl.org/ modeling, and other domains [15, 10], we investigate the effects of applying learned transformations to *in silico* sensitivity screens of cellular models. We modulate gene expression patterns by small molecule representations through learned shifting, scaling, and affine transformations. We compare the predictive performance of these models to baselines such as a model that concatenates raw features. Finally, we investigate conditioning parameters and perform controls such as removing small molecule structural information to understand model dependence on these features.

## 2 Related Work

We expect that combining structured representations of biological information with drug information will improve prediction performance of cellular viability. By designing models with stronger relational inductive biases, we can investigate methods of integrating learned representations [1, 7]. Traditional machine learning methods gain performance by integrating multiple independent datasets and annotated biological pathway information [8]. Multi-drug models significantly outperform single-drug models [13]. In the precision oncology domain, deep learning has been applied to cancer classification, prediction of drug response and drug synergy, drug repositioning, and drug discovery [3]. Chang et al. [2] predict drug IC50s by feature extraction of 787 cell line mutational signatures and 244 small molecule fingerprints, which they integrate by “virtual docking”, or concatenation, followed by additional convolutional layers. Their investigation compares learned data type representations, but does not assess methods of integration. Additionally, the validation datasets include virtually all cell lines and drugs already available in the training datasets. Chiu et al. [4] pre-train mutation and expression encoders on TCGA data by means of an autoencoder. The authors then train the initialized encoders on CCLE data. The network concatenates extracted cellular features as input into an IC50 prediction network. By contrast, in this work, we assess the impact of integrating learned representations for cellular features with small molecule features for the prediction of individual cell line sensitivity to oncogenic therapies.

## 3 Methods

We leveraged well-characterized cellular cancer models from large-scale drug screens. The Cancer Therapeutics Response Portal (CTRP) reports the percent viability of 830 cancer cell lines in response to 545 small molecules across a range of dosages (µmol) after data processing [16]. We generated 512-bit RDKit Morgan fingerprints from small molecule SMILES and concatenated them with compound dosages. For molecular characterization of cellular models we queried the Cancer Cell Line Encyclopedia (CCLE) [11]. The Dependency Map Portal provided access to uniformly processed versions of both datasets [9]. We include standardized RNA-seq Transcripts Per Million (TPM) data from the CCLE for cellular state features. To accelerate training time and limit potential overfitting, we restrict cellular features to the L1000 gene set. The final dataset consists of approximately 5.7 million labeled examples, which we randomly split into into 5 cross-validation folds, stratified by cell line, to asses model generalizability to unseen tumor types (Figure 1).

**Figure 1:**
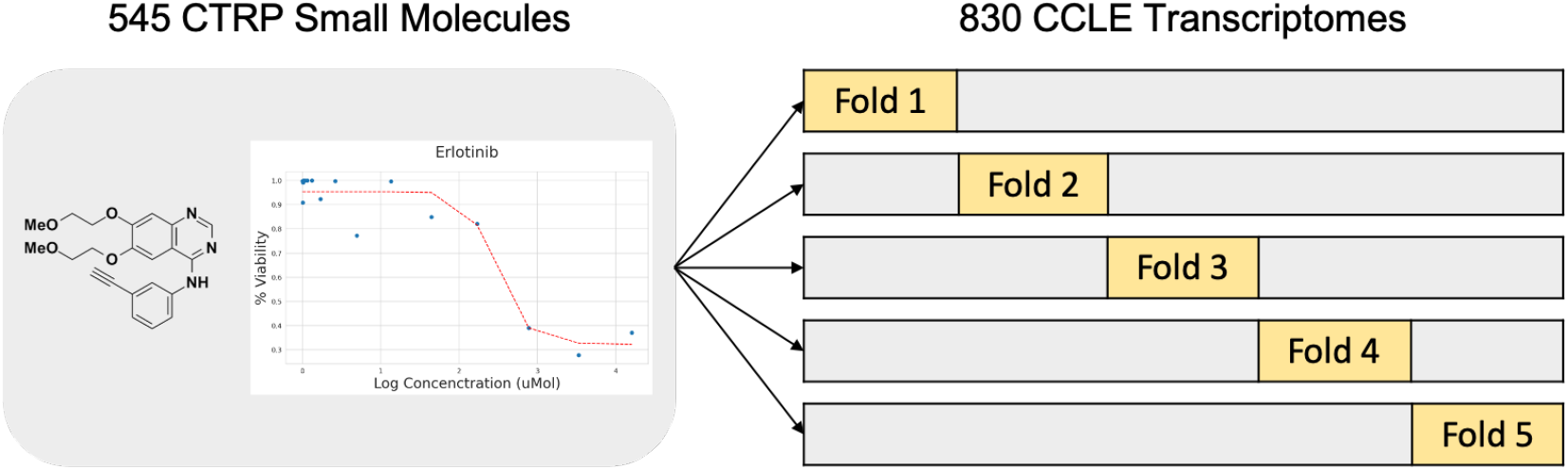
Dataset layout. 545 small molecule drugs tested across a range of dosages in up to 830 unique cancer cell lines comprise 5,767,552 average percent cellular viability measurements. Cell lines were split into 5 folds and models were trained and validated on L1000 gene expression values, conditioned by small molecule structures and dosages, for prediction of percent cellular viability.

We formulate the cellular drug sensitivity prediction task as the conditional model:

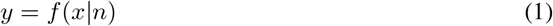

where *y* is cellular sensitivity, *x* is a matrix of standardized RNA-seq TPM values by cell line, and *n* is a matrix of drug dosages and chemical structure fingerprints. Phenotypic response–percent viability–of a cell line depends upon how cells respond to a given small molecule perturbation. As such, we hypothesize that better integration of cell state features with drug structure and dose features will result in more accurate predictions of cellular response.

One means of conditional biasing is simple concatenation of a neural network’s input features. Alternatively, hierarchical representations based on gene expression could instead be combined with learned small molecule feature representations in downstream layers such that a network learns conditional transformation parameters. In our experiments, we assess models that conditionally modulate gene expression features by small molecule features through several different learned transformations.

Under Perez et al. [15]’s general formulation of this approach, termed the feature-wise linear modulation (FiLM) of input features, an affine transformation of inputs by conditional information captures cases of input feature scaling and/or shifting (Figure 2). Model parameters may be learned by functions, *g* and *h*, dependent on conditional information, *n*:

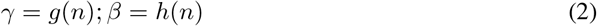

**Figure 2:**
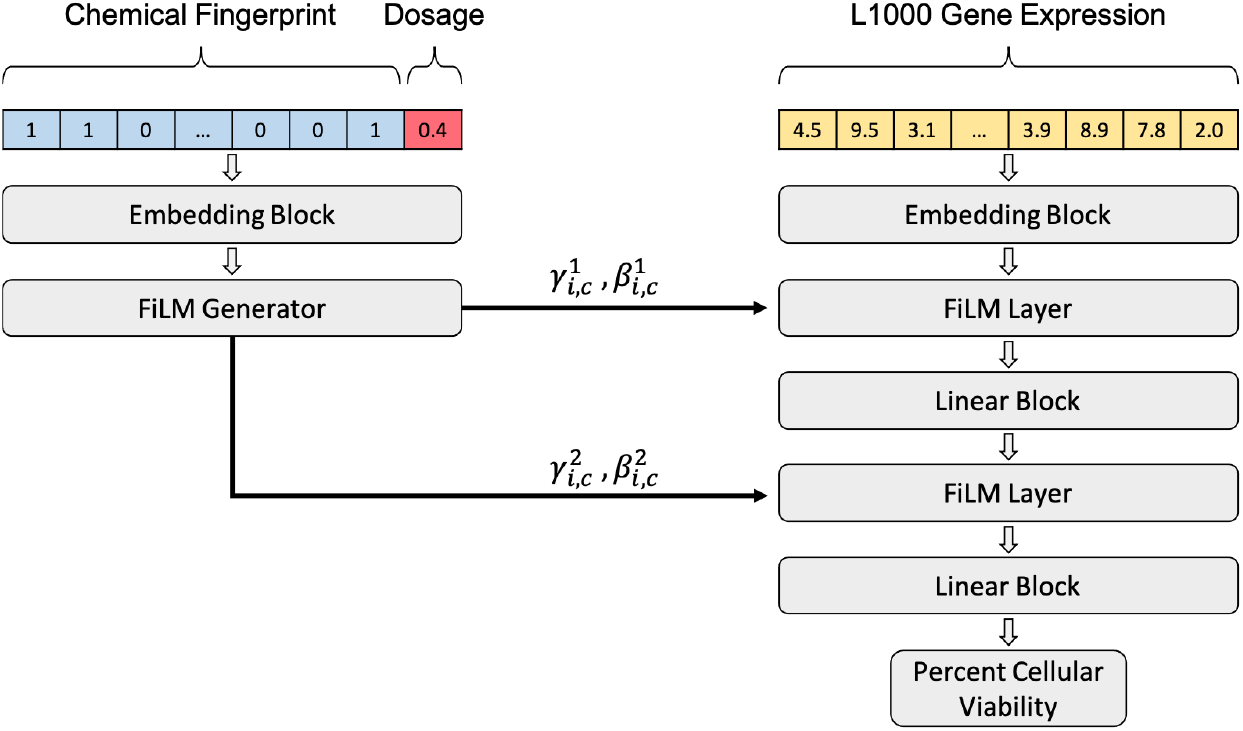
Architecture of conditional cellular sensitivity model. L1000 gene expression representations (right) are conditioned by small molecule representations (left) using an architecture based on FiLM. Learned parameters are applied element-wise for modulation of input features to predict cellular percent viability as a function of cellular gene expression conditioned on drug chemical structure. *γ* and *β* subscripts refer to the *i^th^* input’s *c^th^* feature map.

Learned parameters then modulate intermediate features of a neural network by element-wise trans-formation:

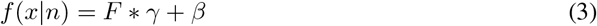

where *F* denotes the activations of a neural network at a given layer.

## 4 Experiments and Results

We assess several means to integrate learned cellular and learned drug structure information in a single model. As a baseline, we compare the “vanilla” approach of raw feature concatenation to three methods of learned feature conditioning [15]. We compare modulation methods by the average maximum coefficient of determination (*R*^2^) of models with similar architecture and training regimens, trained and validated on the same data folds (Table 1). We find that modulation of gene expression features by learned molecular representations (*R*^2^ = 0.7190) outperforms raw feature concatenation (*R*^2^ = 0.6629). Small molecule representation integration by scaling (*R*^2^ = 0.7105), shifting (*R*^2^ = 0.7052), and linear modulation (*R*^2^ = 0.7190) perform similarly, demonstrating that a learned affine transformation captures conditioning of inputs in this domain better than raw feature concatenation. Furthermore, the FiLM model captures the true distribution of percent cellular viability better than the vanilla model. The vanilla model fails to capture low percent cellular viability values (Figure S1). As a consequence of this formulation, the conditioning model architecture implicitly learns valuable “task representations”, i.e. small molecule representations that influence gene expression [10]. Decomposition of learned parameters by t-SNE qualitatively reveals that small molecule dosage dominates the conditioning layer, which is unsurprising given the common sigmoidal relationship between drug response and small molecule concentration (Figure 3) [12]. To quantify the extent to which the dosage variable drives the conditioning, we replaced small molecule structural features with unique but structurally uninformative identifiers of the same length in the concatenation model. This straw model that lacks chemical structure fails catastrophically (*R*^2^ = 0.3327) (Table 1) [5, 6]. These results provide evidence that small molecule structural features are essential in the formulation of our prediction task. Our code is available at https://github.com/keiserlab/film-gex.

**Table 1:** Average coefficients of determination under 5-fold cross validation and identical training regimens for models predicting drug-conditioned cellular viabilities from gene expression. FiLM models outperform scaling or biasing modulation of gene expression values. A straw model trained on data absent of structural features (Drug ID only) fails to explain comparable amounts of percent cellular sensitivity variance.

| Model | $R^2 \pm \text{std. error}$ |
| --- | --- |
| Drug ID only | $0.3327 \pm 0.0977$ |
| Vanilla | $0.6629 \pm 0.0075$ |
| Shift | $0.7052 \pm 0.0144$ |
| Scale | $0.7105 \pm 0.0104$ |
| FiLM | **$0.7190 \pm 0.0098$** |

**Figure 3:**
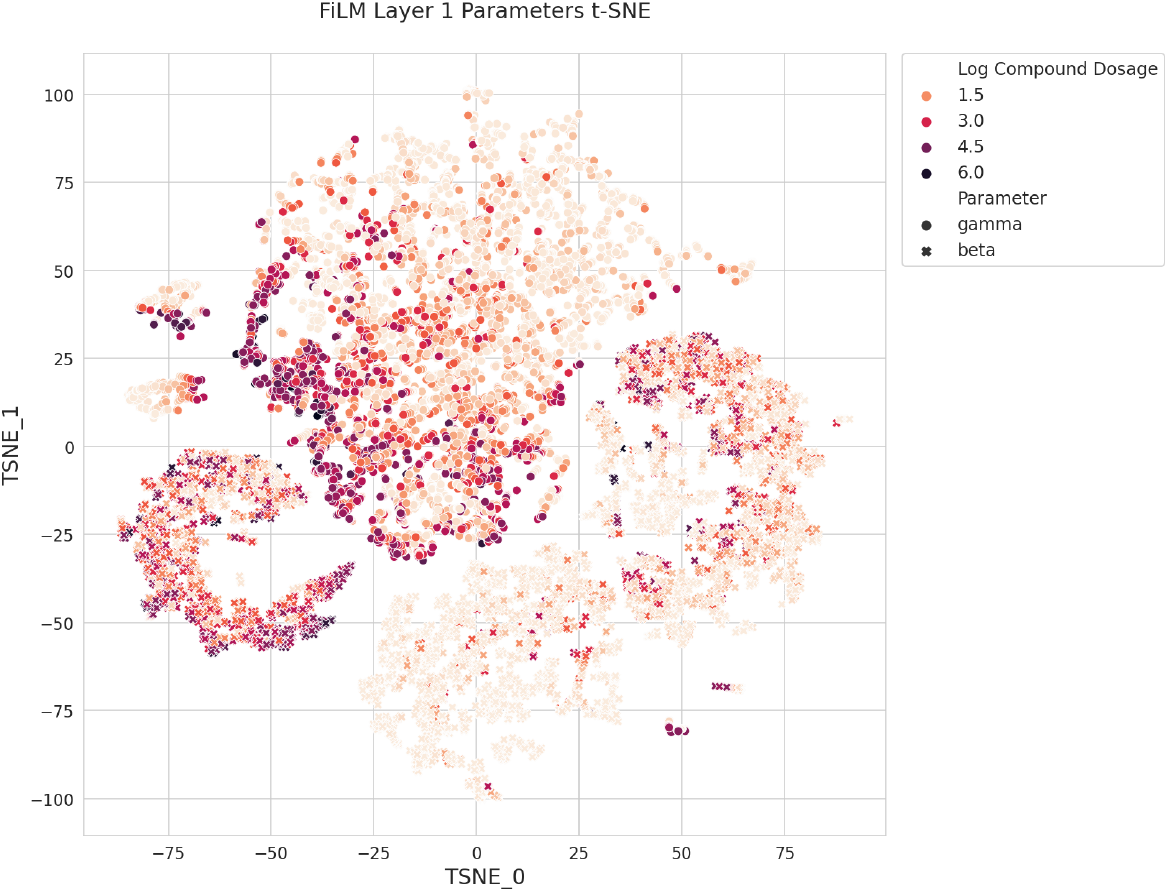
Parameters cluster by small molecule dosage in t-SNE of 32 dimensional FiLM parameters, supporting the substantial role of drug dosage in the model’s modulation.

## 5 Conclusion and Future Directions

We set out to evaluate methods of integrating diverse information by formulating cellular response to a drug perturbation as a conditional model. We compare a conventional neural network model architecture to a set of models with stronger inductive biases. We evaluate conditional modulation of gene expression by learned small molecule representations through shifting, scaling, and affine transformations. Our results show that an explicit conditional model formulation, regardless of applied feature-wise modulation, enhances prediction of cellular sensitivity from diverse data types. The overall success of this task is difficult to compare across reports due to model, data, and outcome subtleties, which we recognize as a benchmarking hurdle. In future work we would be interested to consider cellular features beyond L1000’s 978 gene expression values and simple small molecule fingerprints for better performance. Additionally, more direct comparisons to previous work can be made by restricting the task to prediction of IC50, albeit at substantial detriment to dataset size. Another avenue of pursuit within this conditional framework is *in silico* evaluation of drug synergy/antagonism, in which combinations of task representations modulate cellular features at intermediate network layers. In general, it appears promising to frame biological experimentation as a conditional model, in which effect on a baseline biological state depends upon addition or removal of specific perturbations, whose learned feature representations modulate the model.

## 6 Supplementary Materials

**Figure S1:**
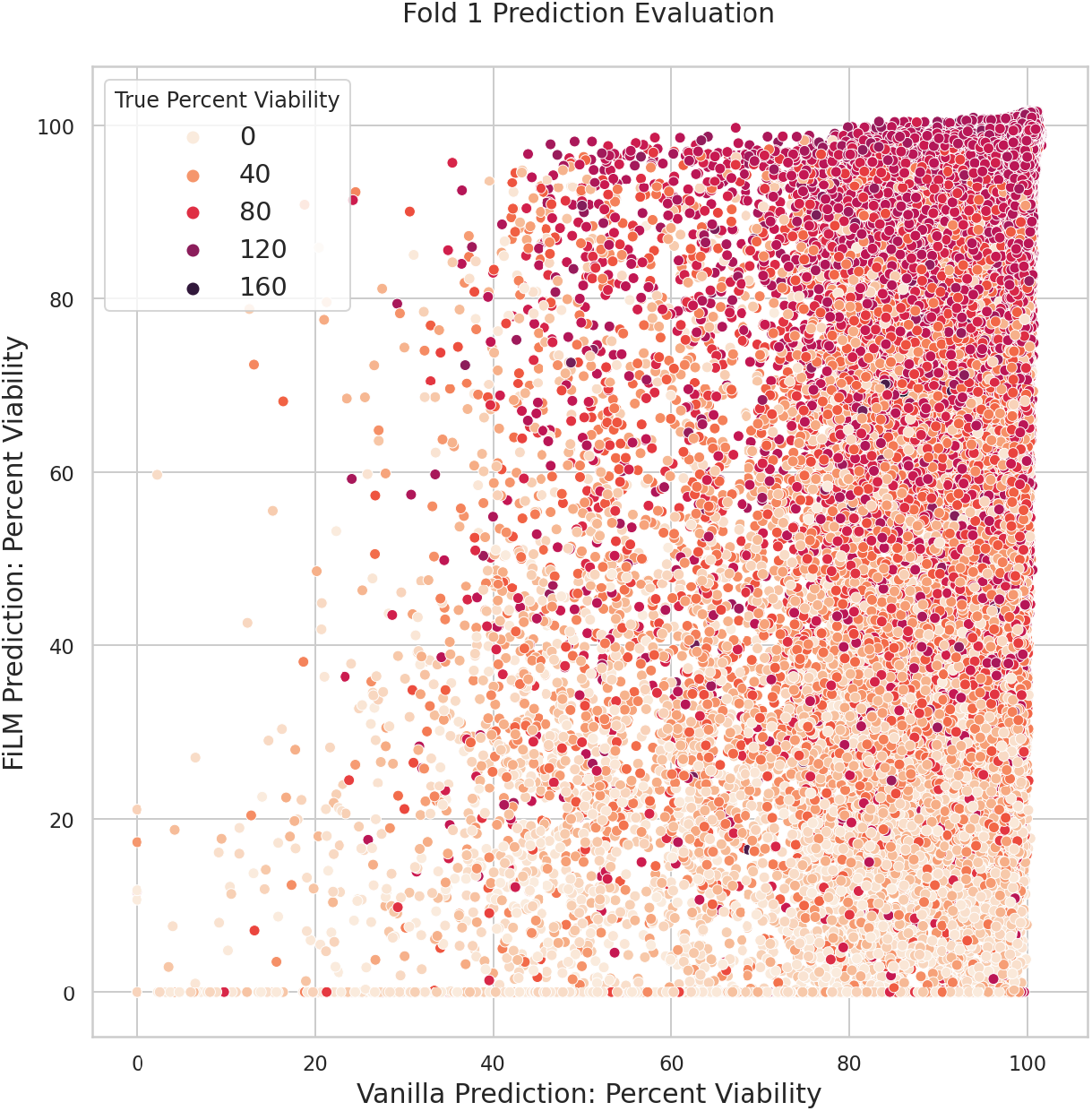
The FiLM model captures the low-end distribution of true percent cellular viability better than the vanilla model.

